# SIRPγ modulates effector differentiation of human CD8 T Cells under suboptimal TCR stimulation: implications for immune homeostasis and autoimmunity

**DOI:** 10.1101/2025.07.09.663913

**Authors:** Megan Morse, Xanthie Rodriguez, Erika DeLaRosa, Sierra Rodriguez, Juma Shanil, Sushmita Sinha

## Abstract

**Background:** Aberrant CD8 T-cell differentiation contributes to the pathogenesis of autoimmune diseases, and immune-mediated tissue damage. However, the molecular mechanisms that prevent premature effector T cell programming in humans remain incompletely defined. Signal regulatory protein gamma (SIRPγ) is selectively expressed on T-cells in the human immune system. Notably, genetic variants associated with reduced SIRPγ expression have been linked to increased risk of immune-mediated diseases, including type 1 diabetes and multiple sclerosis, but the contribution of SIRPγ to CD8 T-cell dysregulation in these contexts remains unclear.

**Objective:** To determine how inter-individual variation in SIRPγ expression influences immune homeostasis and CD8 T-cell effector programming.

**Methods:** Peripheral blood CD8 T-cells from healthy donors were analyzed for SIRPγ expression and associated differentiation phenotypes. Naïve CD8 T-cells were purified and subjected to siRNA-mediated knockdown of *SIRPG*, followed by suboptimal TCR stimulation. Differentiation status, transcription factor expression, and effector cytokine production were measured using flow cytometry. CD47 blockade was used to assess ligand dependency.

**Results:** Low SIRPγ expression on CD8 T-cells was associated with increased frequencies of CD27⁻CD45RO⁺ effector-like and CD27⁻CD45RO⁻ terminally differentiated CD8 T-cells. SIRPG knockdown induced effector-like differentiation, with increased CD45RO and T-bet expression and elevated TNF-α, IFN-γ, and Granzyme B production. This effect was not recapitulated by CD47 blockade, suggesting a CD47-independent regulatory mechanism.

**Conclusion:** SIRPγ serves as a negative regulator of CD8 T-cell effector differentiation under suboptimal stimulation. Inter-individual variation in its expression may influence susceptibility to immune dysregulation, positioning it as a potential biomarker and therapeutic target.

## Introduction

During the COVID-19 pandemic, we have seen the interindividual variability in immune responses to SARS-CoV-2. This variability highlights a key long-standing and unresolved question in immunology: why do some individuals mount a balanced immune response that eliminates infection with no harm to host cells, whereas others have exaggerated immune responses that can cause significant tissue injury and may precipitate autoimmunity(1–9)? Despite extensive research, the host-intrinsic factors that fine-tune T cell responses and immune homeostasis remain incompletely understood. Uncovering these mechanisms is critical for addressing not only infection but also autoimmunity and cancer, where immune balance is often disrupted.

T cells play a critical role in shaping a balanced immune response to foreign antigens and self-antigens. They do so by directly recognizing molecules expressed on the cell surface and secreting factors that drive or dampen local inflammatory responses. One such molecule is signal regulatory protein gamma (SIRPγ), an immunomodulatory protein that is uniquely expressed on the cell surface of human T cells(10, 11). Variants in the *SIRPG* gene have been associated with type 1 diabetes(12–15), relapsing remitting multiple sclerosis, and maintaining long-term vaccine responses(16). We found that humans express varying levels of SIRPγ on their T cells, and that T cells with less SIRPγ surface expression exhibit heightened effector status(17). Importantly our prior studies showed that SIRPγ^low^ T cells are significantly enhanced in individuals with autoimmune diseases such as Type 1 diabetes and relapsing remitting multiple sclerosis (RRMS) (18). This suggests a potential role for SIRPγ as a regulatory checkpoint whose dysfunction contributes to human autoimmunity. Since the publication of our original findings, a slew of papers has emerged describing the role of SIRPγ in chronic immune responses(19), lung squamous cell carcinoma(20), cancer stem-like cells(21), and Type 1 diabetes(15, 22). However, its specific role in the immune system, particularly naïve CD8 T cells, remains unidentified.

Naïve CD8⁺ T cells, upon encountering antigen, undergo extensive phenotypic and functional changes, transitioning through central memory (CM), effector memory (EM), and terminally differentiated effector (TEMRA) stages. must be tightly regulated to prevent inappropriate effector commitment, especially under low-affinity or suboptimal stimulation. Previous studies, including our own, have shown that *SIRPG* expression is elevated in naïve and CM CD8⁺ T cells, with expression progressively declining as cells transition to EM and TEMRA states(17, 19). While this trend suggests a link between SIRPγ and effector stages, it has not been clear whether SIRPγ plays an active role in modulating differentiation or is simply a marker of T cell maturity.

In this study, we address this knowledge gap by investigating the functional role of SIRPγ in regulating human CD8⁺ T cell differentiation. We demonstrate that inter-individual variability in SIRPγ expression-both naturally occurring and experimentally induced via siRNA-profoundly influences the balance between naïve and effector T cell subsets. Specifically, SIRPG knockdown in naïve human CD8⁺ T cells leads to premature effector-like differentiation under suboptimal TCR stimulation. characterized by CD27^low^ CD45RO^+^ phenotype and elevated expression of effector cytokines and transcription factors. Notably, these effects were independent of CD47, the known SIRPγ ligand, suggesting a distinct signaling role for SIRPγ in regulating differentiation thresholds.

By identifying SIRPγ as a molecular brake that prevents premature effector programming, our findings reveal a mechanism by which human T cells resist inappropriate activation under weak antigenic signals. These insights are particularly relevant to contexts such as autoimmunity, chronic infections, and vaccine responsiveness, where immune dysregulation can lead to tissue damage or immune exhaustion. Moreover, given the lack of a murine ortholog, this work underscores a uniquely human regulatory axis in T cell biology—one that may offer novel therapeutic opportunities for modulating effector responses in disease.

## Methods

### Study participants

De-identified leukoreduction buffy coat samples were collected from 44 healthy donors (HDs) at Gulf Coast Regional Blood Center, Houston, TX. All studies were conducted in compliance with the Declaration of Helsinki and approved by the Texas Woman’s University Institutional Review Board (IRB). The mean age of the HDs was 40 ± 18 years, with a gender distribution of 21 males to 19 females. All the reagents used in the study are summarized in Table 1.

**Table 1:** List of reagents used in the study.

| Reagents | Manufacturer | Product # |
| --- | --- | --- |
| Lymphoprep | Stemcell Technologies | 7851 |
| EasySep Human Naïve CD8+ T cell Isolation Kit II | Stemcell Technologies | 17968 |
| Accell Non-targeting Control Pool | Horizon Dharmacon | D-001910-10-20 |
| Accell Human SIRPG siRNA | Horizon Dharmacon | E-013337-00-0005 |
| Anti-Human CD3 | Tonbo Biosciences | 40-0037 |
| Anti-Human CD28 | Biolegend | B314394 |
| Phosphate-buffered saline | Corning | 21040CV |
| 1% paraformaldehyde | Tonbo Biosciences | TNB-8222 |
| Foxp3/Transcription Factor Staining Buffer Kit | Tonbo Biosciences | TNB-0607 |
| IgG1 Isotype Control | R&D systems | MAB002 |
| Anti-hCD47 | R&D systems | MAB4670 |
| APC-700-CD3 | BD bioscience | 3228158 |
| PE-Cy7-CD4 | BD bioscience | 2182173 |
| BV785-CD8 | Biolegend | 344740 |
| PE-SIRPγ | Biolegend | 336606 |
| Alexa Fluor 488-CD27 | Biolegend | 393204 |
| BV650-CD45RO | Biolegend | 304232 |
| PE/Cy7- CD47 | Biolegend | 323114 |
| APC-T-bet | Biolegend | 644814 |
| PE/Cy7- TNF-α | Biolegend | 502930 |
| Alexa Fluor 700- IFN-γ | BD biosciences | 2144428 |
| BV421- Granzyme B | BD biosciences | 3135489 |

### Cell preparation, knock-down and activation

Peripheral blood mononuclear cells (PBMCs) were isolated from buffy coats using a density gradient with Lymphoprep (Stemcell Technologies, Vancouver, BC). SIRPγ^high^ and SIRPγ^low^ donors were identified as described earlier(17). PBMC samples from SIRPγ^high^ and SIRPγ^low^ donors were used for determining the frequencies of naïve, central memory (CM), effector memory (EM) and terminal effectors (TEMRA) in SIRPγ^high^ vs SIRPγ^low^ donors. Naïve CD8 T cells were then purified from the PBMC samples from SIRPγ^high^ donors using a CD8 Naïve T Cell Sorting Kit from Stemcell Technologies, following the manufacturer’s instructions. The purity of the naïve CD8 T cells was consistently >99%. The purified cells were cultured with either scrambled scrambled control siRNA or a SMARTpool siRNA targeting *SIRPG* (Horizon Discovery, Cambridge, UK), with knock-down performed according to the manufacturer’s guidelines. For activation, cells were treated with suboptimal concentrations of anti-CD3 (Tonbo Biosciences, San Diego, CA) or 48 hours as described previously. We chose sub-optimal activation of naïve CD8 T cells because our previous research demonstrated that SIRPγ^low^ CD8 T cells have lower activation thresholds(17). Following incubation, the plates were centrifuged, and the cells were used for flow cytometry staining.

### Flow Cytometry staining

Fluorescently conjugated antibodies used in the study are listed in Table 1. Cell samples were washed with phosphate-buffered saline (PBS, Mediatech Cellgro) and then stained with the appropriate fluorescently labeled antibodies. After staining, cells were resuspended in 1% paraformaldehyde (Tonbo Biosciences, San Diego, CA). Flow cytometry data were acquired using a CytoFLEX Flow Cytometer (Beckman Coulter, Brea, CA) and analyzed with FlowJo software (BD Biosciences, San Jose, CA). Intracellular flow cytometry staining was performed to detect IFNγ, TNFα, Granzyme B and T-bet. Briefly cells were fixed and permeabilized (Tonbo Biosciences, San Diego, CA) before intracellular staining.

### CD47 neutralization assay

Sorted naïve CD8 T cells were plated in 96-well plates, and control-IgG or anti-CD47 neutralizing antibody was added(23). Cells were activated after 2 hours with suboptimal concentrations of anti-CD3. Following incubation, the plates were centrifuged, and the cells were used for flow cytometry staining.

## Results

### SIRPγ expression varies between individuals and stratifies CD8⁺ T cell differentiation states

We have shown that SIRPγ expression on CD8⁺ T cells varies significantly among individuals(17). While most individuals exhibit high levels of SIRPγ (Fig. 1A), a subset (carrying the SNP rs2281808) displays markedly low expression (Fig. 1C). Beyond natural variability, SIRPγ expression is also influenced by T cell differentiation. Specifically, SIRPγ is robustly expressed on naïve and central memory T cells but decreases as T cells differentiate into effector and terminally differentiated subsets (SI). This pattern suggests that SIRPγ may play a role in regulating T cell differentiation. To investigate this further, we compared the differentiation profiles of CD8⁺ T cells from individuals with high and low SIRPγ expression.

**Figure 1.**
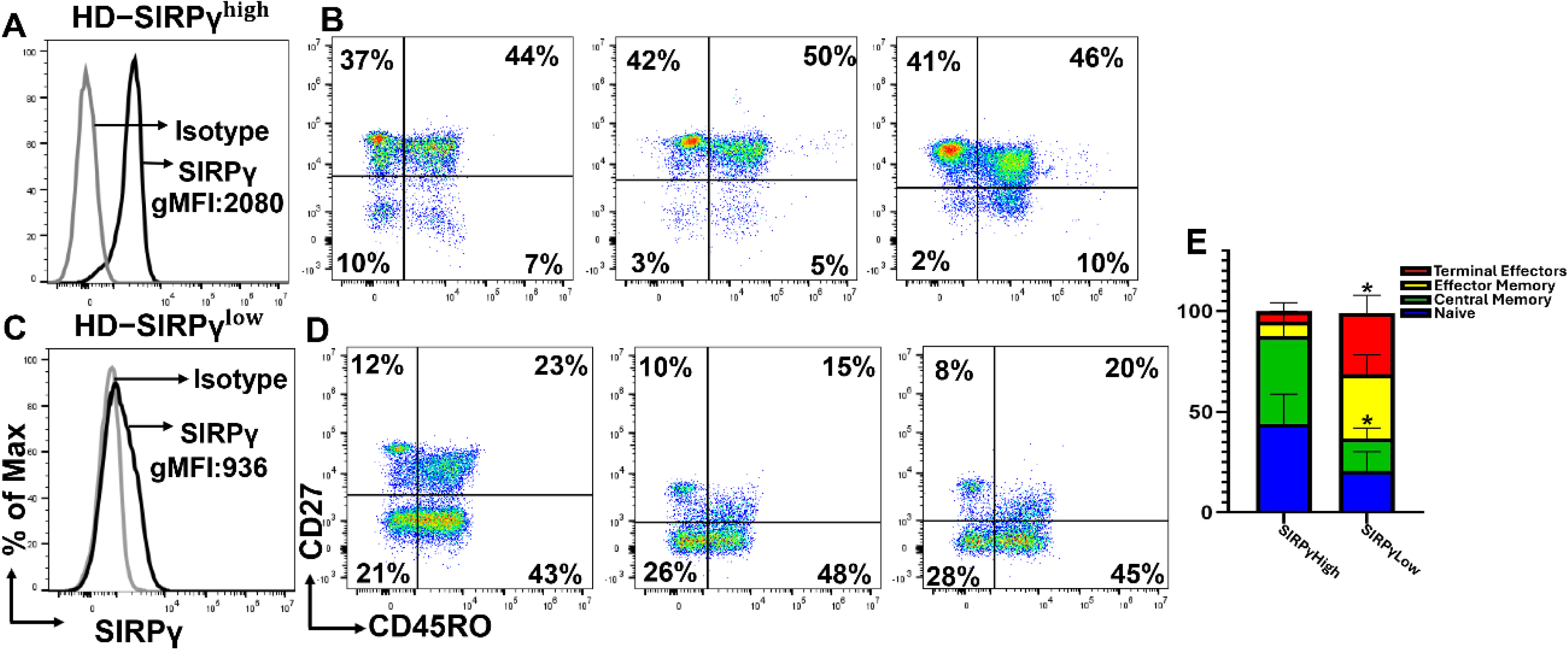
SIRPγ^low^ carriers have significantly increased frequencies of effectors and terminal effectors in peripheral blood. Peripheral blood mononuclear cells (PBMCs) were isolated from 50 Healthy donors (HDs) and stained with fluorescently conjugated antibodies. SIRPγ^high^ vs. SIRPγ^low^ carriers were identified based on a previously described strategy. Representative histograms showing SIRPγ expression on bulk CD8 T cells from a SIRPγ^high^ and a SIRPγ^low^ donor are shown in panels **A &C** respectively. Dot plots showing the distribution of CD8 T cell subsets from three independent SIRPγ^high^ and SIRPγ^low^ donors are shown in panels B and D, respectively. **E** Cumulative data show that SIRPγ^low^ individuals exhibit a significantly higher frequency of effector-like (CD45RO⁺CD27⁻) and terminally differentiated (CD45RO⁻CD27⁻) CD8 T cells compared to SIRPγ^high^ donors. SIRPγ^high^ individuals predominantly retained naïve and central memory-like populations, suggesting a more balanced differentiation state. Statistical analysis was done using an unpaired t-test, *p < 0.001.

Our findings revealed that individuals with low SIRPγ expression have a significantly higher proportion of effector-like (CD45RO⁺CD27⁻, SIRPγ^high^ vs. SIRPγ^low^; Mean ± SD, 6.7 ± 3.7 vs. 32 ± 9.6; p<0.00001) and terminally differentiated T cells (CD45RO⁻CD27⁻ SIRPγ^high^ vs.

SIRPγ^low^; Mean ± SD, 4.7 ± 4 vs. 30 ± 8.7, p<0.00001) (Fig 1D &E), while those with high SIRPγ expression predominantly exhibit naïve and central memory-like cells (Fig 1 B & E). These observations suggest that low SIRPγ expression may skew T cell differentiation toward more differentiated, effector-like states, potentially disrupting immune homeostasis in SIRPγ^low^ individuals. This variability provides a unique opportunity to explore how differential SIRPγ expression impacts T cell differentiation, particularly the transition between naïve, memory, effector, and terminally differentiated states.

### SIRPγ knockdown promotes effector-like differentiation of naïve human CD8⁺ T cells under suboptimal stimulation

To determine whether *SIRPG* knockdown drives effector-like differentiation in naïve human CD8⁺ T cells, we first established an siRNA-mediated knockdown (KD) model in primary human naïve CD8⁺ T cells. Naïve CD8⁺ T cells were isolated at >99% purity from peripheral blood of individuals with high endogenous SIRPγ expression (SIRPγ^high^ donors) and transfected with either scrambled control siRNA or a SMARTpool siRNA targeting *SIRPG*. Forty-eight hours post-transfection, flow cytometry analysis confirmed a substantial reduction in surface SIRPγ expression in the knockdown condition relative to control (Figure 2A, B), demonstrating effective silencing of the target protein. This system enabled us to examine the downstream impact of SIRPγ loss on T cell differentiation under defined stimulatory conditions.

**Figure 2.**
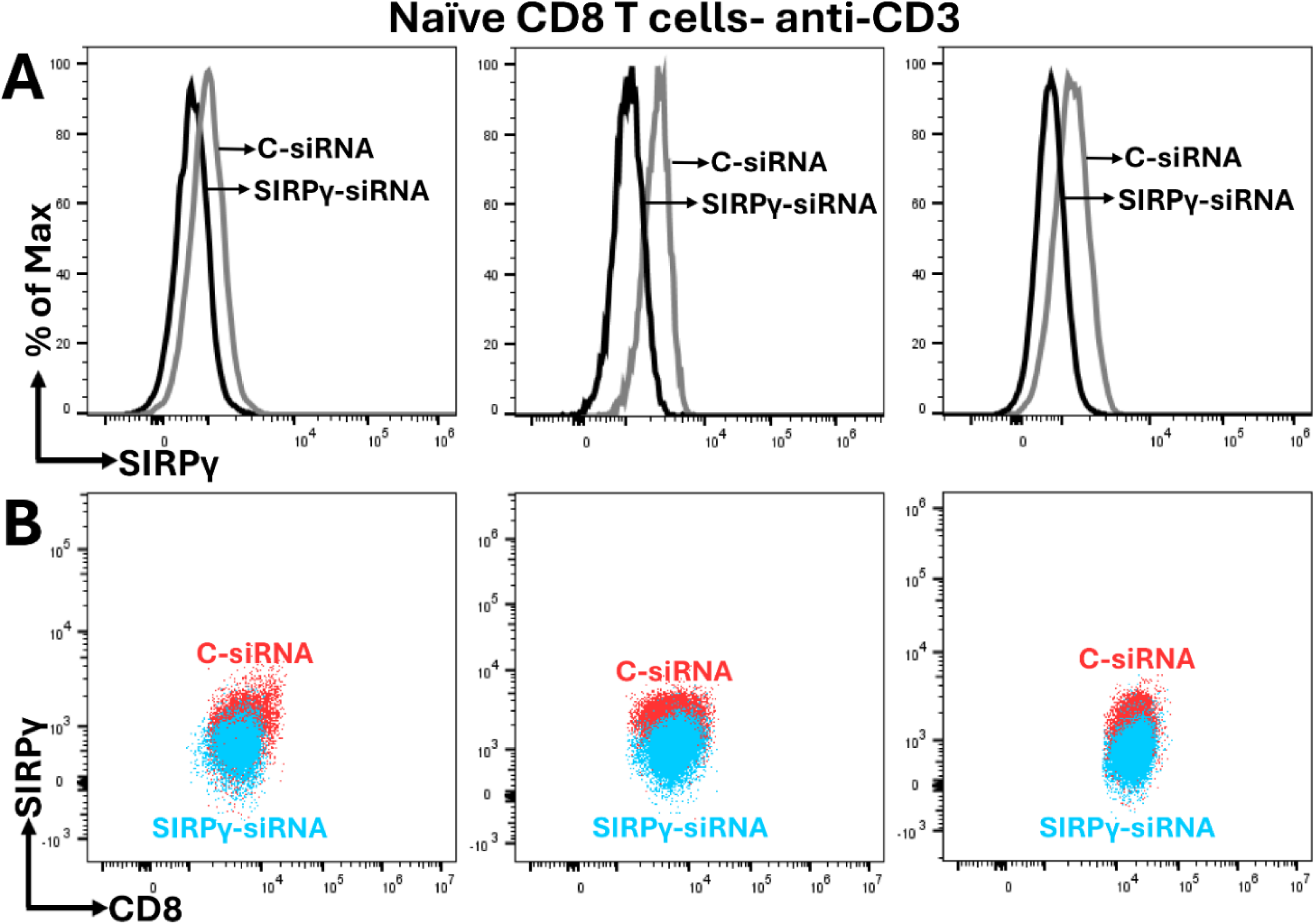
siRNA mediated knockdown (KD) of SIRPγ in naïve human CD8 T cells. Naïve human CD8 T cells were purified from peripheral blood mononuclear cells (PBMCs) of SIRPγ^high^ healthy donors using Stem Cell kit. Cells were transfected with either control scrambled siRNA or a SMARTpool Accell siRNA targeting *SIRPG*, following the manufacturer’s instructions. (A, B) Representative flow cytometry plots showing SIRPγ surface expression in naïve CD8⁺ T cells 48 hours post-transfection. SIRPγ knockdown resulted in a marked reduction in surface expression compared to the scrambled siRNA control.

Following KD, cells were either left unstimulated or subjected to suboptimal stimulation with anti-CD3 antibodies for 72hrs and differentiation was analyzed by flow cytometry. Effector-like cells were defined by the surface phenotype CD27^low^CD45RO^+^. Upon stimulation with suboptimal anti-CD3, *SIRPG* knockdown substantially increased the proportion of CD27^low^CD45RO^+^ve effector-like cells across all donors analyzed. Control samples exhibited low levels of differentiation (ranging from 0.3% to 5%; mean ± SD, 2.5±1.7; Fig. 3A&C), whereas *SIRPG*-deficient cells demonstrated a striking rise in effector-like cells, ranging from 24% to 60%, mean ± SD, 38±11; Fig. 3B & C) under identical conditions. These results demonstrate that *SIRPG* knockdown markedly enhances the capacity of naïve CD8⁺ T cells to adopt an effector-like phenotype, particularly under suboptimal TCR stimulation. This suggests that SIRPγ acts as a regulatory checkpoint, restraining premature effector differentiation when TCR signals are weak.

**Figure 3.**
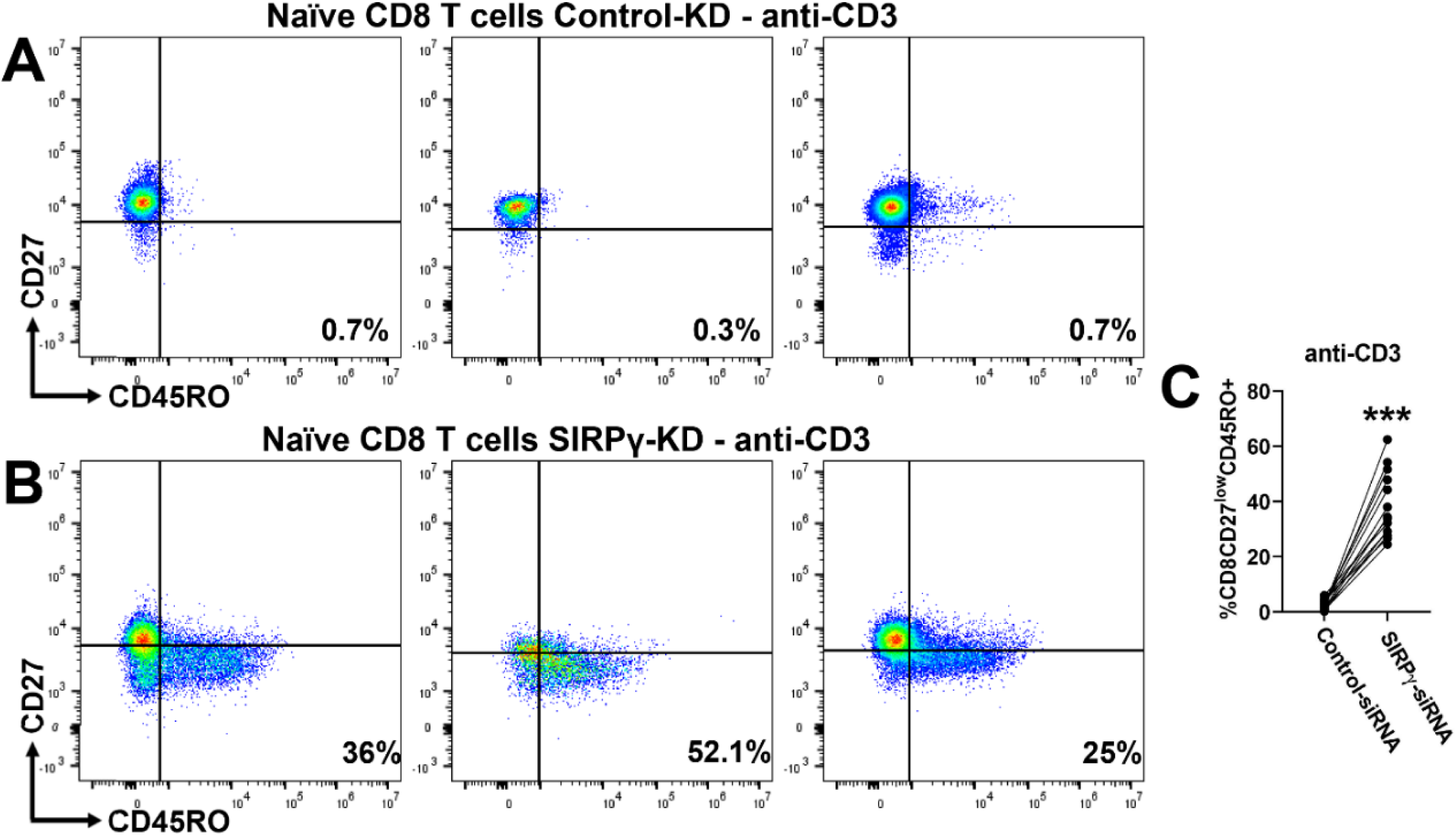
SIRPγ knockdown promotes effector-like differentiation of naïve human CD8 T cells under suboptimal stimulatory conditions. (A, **B)** Representative flow cytometry dot plots showing CD27 and CD45RO expression in naïve CD8 T cells transfected with either control siRNA (**A**) or SIRPG-specific siRNA (**B**) following suboptimal anti-CD3 stimulation. **(C)** Quantification of CD27^low^CD45RO⁺ CD8 T cells from 14 independent SIRPγ^high^ donors reveals a significant increase in effector-like differentiation upon SIRPG knockdown. **(C)**. Statistical analysis was done using a paired t-test, ***p < 0.001.

### Effector-like differentiation of naïve CD8⁺ T Cells following SIRPγ knockdown occurs independently of CD47 interaction

To determine whether the interaction between SIRPγ and its known ligand CD47 is required for effector-like differentiation of naïve CD8⁺ T cells, CD47 was neutralized on freshly isolated naïve human CD8⁺ T cells using a blocking antibody. Cells were then subjected to suboptimal TCR stimulation, a condition that promoted CD45RO upregulation following SIRPγ knockdown. Flow cytometric analysis confirmed effective blockade of CD47 on the cell surface (Fig. 4 A vs. B, top panel). However, in contrast to the phenotype observed with SIRPγ knockdown (Fig. 3B), CD47-neutralized naïve CD8⁺ T cells did not upregulate CD45RO upon suboptimal stimulation (Fig 4. B bottom panel & C). These results demonstrate that the effector-like differentiation observed following SIRPγ knockdown is not reproduced by CD47 blockade alone, indicating that SIRPγ’s regulatory role in CD8⁺ T cell differentiation occurs independently of its interaction with CD47. This suggests the involvement of CD47-independent signaling mechanisms downstream of SIRPγ.

**Figure 4.**
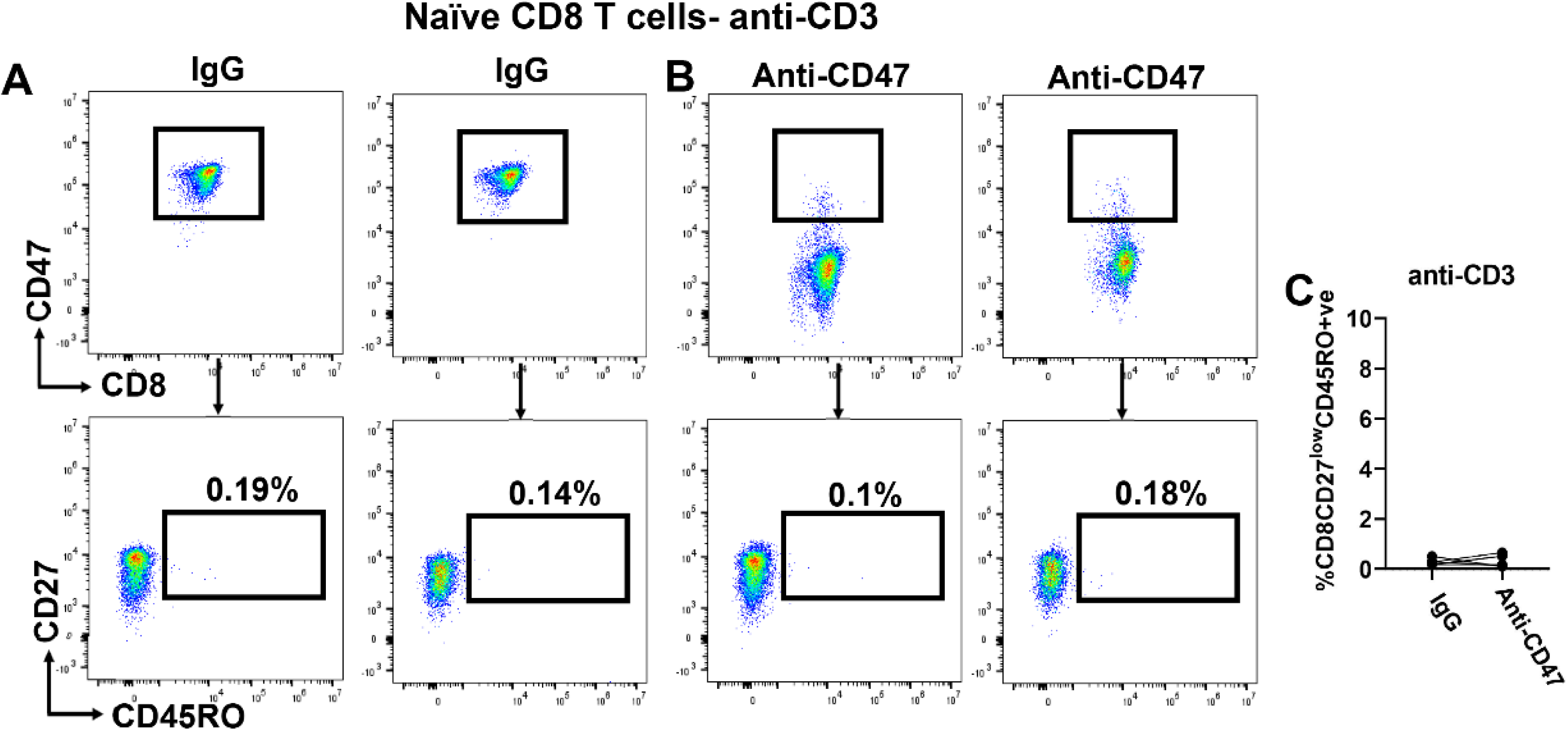
CD47 blockade does not promote effector-like differentiation of naïve CD8⁺ T cells under suboptimal stimulation. Naïve human CD8 T cells from SIRPγ^high^ donors were pretreated with a neutralizing anti-CD47 antibody or isotype control, followed by suboptimal anti-CD3 stimulation for 72 hrs. **(A)** Flow cytometry analysis confirmed effective blockade of CD47. **(B)** Expression of CD45RO was assessed post-stimulation. CD47-neutralized cells did not show an increased CD45RO expression compared to isotype-treated controls. Data are representative of six independent donors. These results indicate that SIRPγ-mediated regulation of CD8⁺ T cell differentiation occurs via a CD47-independent mechanism.

### SIRPγ knockdown enhances T-bet expression in emerging effector CD8 T cells

We have previously shown that T-bet expression correlates inversely with SIRPγ expression in human CD8 T cells, raising the possibility that SIRPγ may restrain effector differentiation by limiting T-bet expression. To directly test this, we analyzed T-bet expression in SIRPγ knockdown cultures following suboptimal TCR stimulation, comparing CD45RO⁺ (effector-like, Fig. 5A&C) and CD45RO⁻ (naïve, Fig. 5A&B) subsets by intracellular flow cytometry. CD45RO⁺ cells showed a clear increase in T-bet expression relative to their naive counterparts in SIRPγ knockdown cultures (SIRPγ KD, CD45RO-vs. CD45RO+ve; Mean ± SD; 1%±0.8% vs. 23%±5; p<0.001; Fig 5C&D). These results suggest that loss of SIRPγ facilitates T-bet upregulation specifically in CD8⁺ T cells that are transitioning into an effector-like state, even under suboptimal stimulation.

**Figure 5.**
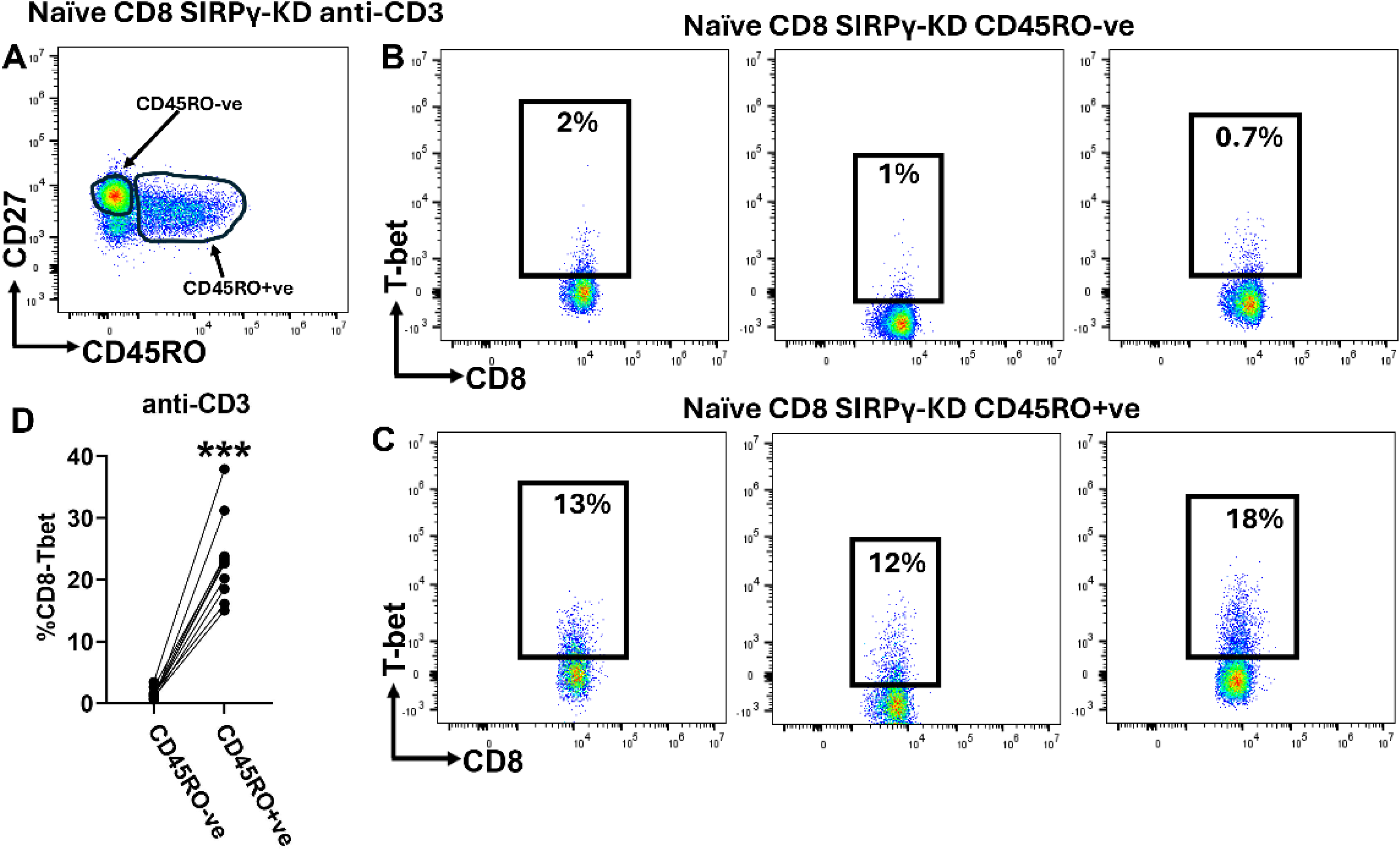
SIRPγ knockdown promotes T-bet expression in CD8 T cell effectors. **(A)** Representative flow cytometry plot showing CD45RO⁻ and CD45RO⁺ CD8 T cell subsets after SIRPγ knockdown. **(B, C)** Intracellular T-bet expression in CD45RO⁻ (**B**) and CD45RO⁺ (**C**) subsets, assessed by flow cytometry in SIRPγ knockdown cultures from three independent donors. CD45RO⁺ cells consistently exhibited significantly higher T-bet expression compared to CD45RO⁻ cells across donors. **(D)** Cumulative data is representative of 10 independent donors. These findings are consistent with our prior observations that T-bet expression inversely correlates with SIRPγ levels in CD8 T cells, supporting a model in which reduced SIRPγ promotes effector differentiation through T-bet–mediated transcriptional programming. Statistical analysis was done using a paired t-test, ***p < 0.001.

### SIRPγ knockdown-induced CD45RO⁺ cells function as bona fide effectors

Given the enhanced expression of T-bet in CD45RO⁺ effector-like populations upon SIRPγ knockdown, we next examined whether these cells functionally behave as effectors. Importantly, we tested this under suboptimal stimulation conditions, using low-dose anti-CD3 to better reveal the impact of SIRPγ loss on early activation thresholds—conditions that typically do not drive robust cytokine production in naïve T cells. Using intracellular cytokine staining and flow cytometry, we compared TNF-α, IFN-γ, and Granzyme B production between CD45RO⁺ and CD45RO⁻ subsets in SIRPγ-deficient cultures after suboptimal anti-CD3 stimulation (Fig. 6 &B). As expected, CD45RO⁻ cells (representing the naive pool) minimal cytokine production under these conditions (Fig. 6A &C), whereas CD45RO⁺ cells consistently showed a significantly higher levels of effector molecules (Fig. 6B & C). We observed a marked increase in TNF-α production in the CD45RO⁺ population across all donors with several donors exhibiting more than a two- to three-fold increase (SIRPγ KD, CD45RO-vs. CD45RO+ve; Mean ± SD; 10%±5% vs. 33%±15; p<0.001; Fig. 6B & C). A similar pattern was observed for IFN-γ, where CD45RO⁺ cells produced significantly more cytokine compared to their CD45RO⁻ counterparts (SIRPγ KD, CD45RO-vs. CD45RO+ve; Mean ± SD; 1%±0.6% vs. 12%±2.7; p<0.001; Fig. 6B&D). In addition, we examined Granzyme B expression as a surrogate for cytotoxic potential. Again, the CD45RO⁺ subset showed enhanced expression as compared to their CD45RO⁻ counterparts, (SIRPγ KD, CD45RO-vs. CD45RO+ve; Mean ± SD; 5%±3% vs. 31%±16; p<0.001; Fig 6B & E). These results demonstrate that CD45RO⁺ cells emerging after SIRPγ knockdown are not only phenotypically distinct but also functionally resemble effector cells, producing key cytokines and cytotoxic molecules. The elevated production of TNF-α, IFN-γ, and Granzyme B confirms that these cells behave as bona fide effectors, supporting the idea that SIRPγ sets an activation threshold that normally limits premature or excessive effector differentiation.

**Figure 6.**
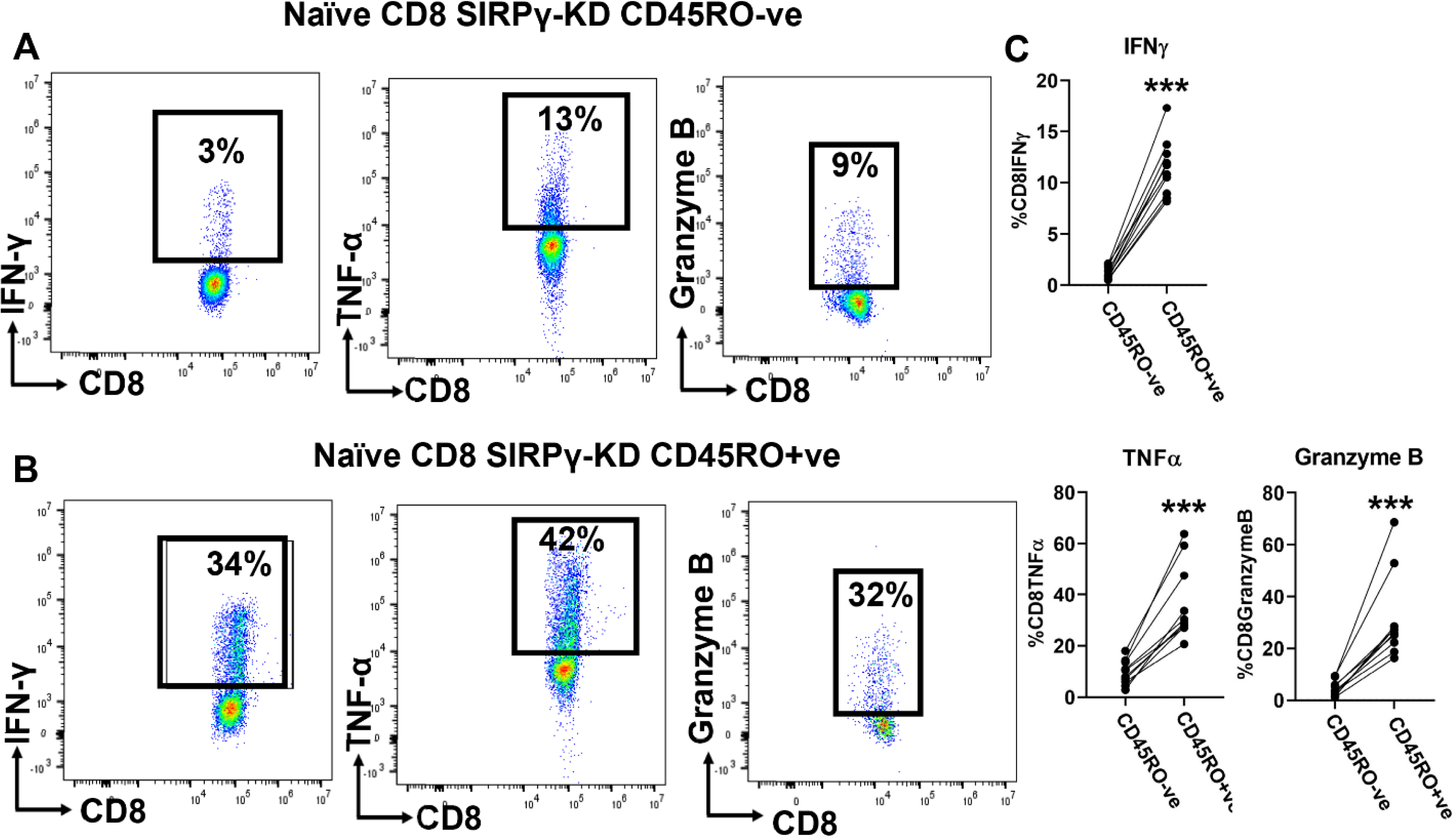
SIRPG knockdown enhances effector cytokines and granzyme production in CD45RO⁺ CD8 T cells. Naïve human CD8 T cells from SIRPγ^high^ donors were subjected to SIRPG knockdown and subsequently analyzed for CD45RO expression as shown in figure 4A. To evaluate effector function, intracellular levels of TNFα, IFNγ, and Granzyme B were measured by flow cytometry in CD45RO^-^ (**A)** and CD45RO^+^ (**B)** subsets. CD45RO⁺ cells produced significantly higher levels of TNFα, IFNγ, and Granzyme B compared to their CD45RO⁻ counterparts, indicating that SIRPG knockdown promotes functional differentiation of CD8⁺ T cells toward an effector phenotype. **(C)** Cumulative data from 10 independent SIRPγ^high^ donors. Statistical analysis was done using a paired t-test, ***p < 0.001.

## Discussion

SIRPγ is a member of the signal-regulatory protein (SIRP) family that is selectively expressed on human T cells in the immune system(11). While its expression patterns across CD8⁺ T cell subsets have been described previously by us and others, its functional role in regulating human CD8⁺ T cell differentiation has remained poorly understood. Here, we provide new mechanistic and functional insights into how SIRPγ constrains effector differentiation and shapes CD8⁺ T cell fate decisions, particularly under suboptimal stimulatory conditions.

Consistent with our prior observations and those reported by others, we confirm that SIRPγ expression is highest on naïve and central memory CD8⁺ T cells and declines with progressive differentiation into effector and terminally differentiated states. Extending these findings, our current work reveals that this expression pattern is not merely a marker of differentiation status, but that SIRPγ itself plays an active role in modulating CD8⁺ T cell fate. Specifically, we demonstrate that inter-individual variability in SIRPγ expression strongly correlates with the composition of the peripheral CD8⁺ T cell pool. Individuals with abnormally low SIRPγ expression on CD8 T cells have an increased frequency of CD27⁻ CD45RO⁺ effector-like and CD27⁻ CD45RO⁻ terminally differentiated T cells-phenotypes corresponding to more differentiated effector and terminal states. This suggests that low SIRPγ expression may predispose individuals to accelerated or dysregulated T cell differentiation.

To directly test the functional role of SIRPγ in regulating differentiation, we knocked down *SIRPG* in purified naïve CD8⁺ T cells and examined their differentiation in response to low-dose TCR stimulation—a condition mimicking weak or suboptimal antigenic exposure. Knockdown of SIRPG resulted in a robust shift toward an effector-like CD45RO⁺ CD27^low^ phenotype, even in the absence of strong co-stimulation. This phenotype was consistent across multiple donors and underscores a previously unrecognized role for SIRPγ in enforcing a threshold for differentiation under suboptimal stimulatory conditions. These findings suggest that SIRPγ functions as a molecular brake, enforcing a threshold that prevents premature effector differentiation under limited activation conditions.

These findings place SIRPγ among a growing number of molecules that tune effector differentiation through modulation of activation thresholds. Similar to CD5 and NR4A family members, which act as negative regulators of TCR signal strength and prevent hyperactivation(24, 25), SIRPγ may function to prevent low-affinity or sub-threshold TCR signals that would otherwise lead to inappropriate effector programming. In the context of chronic infection or autoimmunity, such a regulatory mechanism would be crucial to maintain immune homeostasis.

Importantly, the CD45RO⁺ effector-like cells emerging from *SIRPG* knockdown cultures exhibited hallmark features of functional effector T cells. They displayed enhanced production of TNF-α, IFN-γ, and Granzyme B upon sub-optimal stimulation, and showed upregulation of the transcription factor T-bet—a key regulator of type 1 effector CD8⁺ T cell differentiation(26, 27). Notably, T-bet expression was restricted to the CD45RO⁺ subset, suggesting that these cells had undergone true effector programming. Thus, SIRPγ-deficient CD8⁺ T cells not only acquire phenotypic markers of differentiation but also gain functional effector capabilities.

Interestingly, the effector-like phenotype induced by SIRPγ knockdown could not be replicated by blocking its known ligand, CD47. Although SIRPγ-CD47 interaction has been proposed to mediate T cell migration(28), CD47 blockade failed to induce CD45RO expression or effector differentiation under suboptimal stimulation. This finding suggests that the regulatory function of SIRPγ in T cells is not solely mediated through CD47-dependent interactions. Our results reveal a CD47-independent role for SIRPγ in intracellular signaling that influences fate decisions. It is possible that SIRPγ may modulate signaling by associating with adaptor proteins or by influencing the spatial organization of signaling complexes at the immunological synapse. These possibilities warrant further investigation.

The interindividual variability in SIRPγ expression, particularly the presence of a naturally occurring SIRPγ^low^ subset, raises important questions about susceptibility to immune dysregulation. It is plausible that SIRPγ^low^ individuals may be predisposed to heightened effector responses, with potential implications for vaccine responsiveness, immunopathology, or even autoimmunity. In this context we have shown that both relapsing remitting multiple sclerosis and type 1 diabetes patients have significantly increased frequencies of SIRPγ^low^ CD8 T cells.

Taken together, our findings define a new role for SIRPγ in human CD8⁺ T cell biology: it acts as a differentiation checkpoint that restricts effector programming under weak stimulatory conditions. This function may be critical for maintaining T cell homeostasis and preventing unnecessary immune activation. Inter-individual differences in SIRPγ expression could therefore have important consequences for susceptibility to immune exhaustion, responsiveness to chronic infections, or propensity for autoimmunity. Notably, our previous work established that SIRPγ^low^ CD8⁺ T cells are enriched in autoimmune conditions including Type 1 Diabetes and RRMS. Coupled with our current findings, this suggests that dysregulated SIRPγ expression may predispose to inappropriate effector T cell differentiation, contributing to the immune pathology observed in these diseases. Moreover, with growing interest in targeting SIRP family members for immunotherapy—particularly in the context of cancer(29)—our data suggest that modulating SIRPγ may represent a novel strategy to fine-tune T cell differentiation and function. Further exploration of the molecular mechanisms and regulatory networks involving SIRPγ will be essential to assess its potential as a therapeutic target in human immunopathology.

## Conclusion

Our findings identify SIRPγ as a critical regulator of human CD8⁺ T cell differentiation. SIRPγ prevents premature effector programming under suboptimal activation conditions. The inter-individual variability in SIRPγ expression, shaped in part by genetic variation, may influence susceptibility to autoimmune and inflammatory diseases. These results highlight SIRPγ as a potential biomarker of immune dysregulation and a novel target for therapeutic modulation of T cell responses in human disease.

## Acknowledgments

Several undergraduate students were trained and participated in various experiments included in this manuscript. We thank Shradda Uppu, Varun Reddy, Aparna Sunny, Victor Diran, Aideth Serrano, Citlaly Torres, Colyn Bessard, Samantha Mendoza for their contributions to buffy coat processing and fluorescent staining. We also extend our gratitude to Dr. Juliet Spencer and Dr. Christopher Brower from TWU for their valuable discussions on methodology and concepts.

## Statement and Declaration

The authors declare that the research was conducted in the absence of any commercial or financial relationships that could be construed as a potential conflict of interest.

## Author contribution

All authors contributed to the study conception and design. Material preparation, data collection and analysis was performed by Megan Morse, Xanthie Rodriguez, Erika Rodriguez, Sierra Rodriguez, Juma Shanil, Sushmita Sinha. First draft of the manuscript was written by Sushmita Sinha and Megan Morse. And all authors commented on the previous versions of the manuscript. All authors read and approved the final manuscript.

## Notes

### Competing Interest Statement

The authors have declared no competing interest.

## References

1. Casadevall A, Pirofski LA. What Is a Host? Attributes of Individual Susceptibility. Infect Immun. 2018;86(2).

2. Jingwu Z, Medaer R, Hashim GA, Chin Y, van den Berg-Loonen E, Raus JC. Myelin basic protein-specific T lymphocytes in multiple sclerosis and controls: precursor frequency, fine specificity, and cytotoxicity. Ann Neurol. 1992;32(3):330–8.

3. Joshi N, Usuku K, Hauser SL. The T-cell response to myelin basic protein in familial multiple sclerosis: diversity of fine specificity, restricting elements, and T-cell receptor usage. Ann Neurol. 1993;34(3):385–93.

4. Lama J, Planelles V. Host factors influencing susceptibility to HIV infection and AIDS progression. Retrovirology. 2007;4:52.

5. Martin R, Jaraquemada D, Flerlage M, Richert J, Whitaker J, Long EO, et al. Fine specificity and HLA restriction of myelin basic protein-specific cytotoxic T cell lines from multiple sclerosis patients and healthy individuals. J Immunol. 1990;145(2):540–8.

6. Sinha S, Crawford MP, Ortega SB, Karandikar NJ. Multiparameter Flow Cytometric Assays to Quantify Effector and Regulatory T-Cell Function in Multiple Sclerosis. Journal of multiple sclerosis. 2015;2(1).

7. Sparling PF. A plethora of host factors that determine the outcome of meningococcal infection. Am J Med. 2002;112(1):72–4.

8. Thair SA, He YD, Hasin-Brumshtein Y, Sakaram S, Pandya R, Toh J, et al. Transcriptomic similarities and differences in host response between SARS-CoV-2 and other viral infections. iScience. 2021;24(1):101947.

9. Tharappel AM, Samrat SK, Li Z, Li H. Targeting Crucial Host Factors of SARS-CoV-2. ACS Infect Dis. 2020;6(11):2844–65.

10. Barclay AN, Van den Berg TK. The interaction between signal regulatory protein alpha (SIRPalpha) and CD47: structure, function, and therapeutic target. Annual review of immunology. 2014;32:25–50.

11. Piccio L, Vermi W, Boles KS, Fuchs A, Strader CA, Facchetti F, et al. Adhesion of human T cells to antigen-presenting cells through SIRPbeta2-CD47 interaction costimulates T-cell proliferation. Blood. 2005;105(6):2421–7.

12. Barrett JC, Clayton DG, Concannon P, Akolkar B, Cooper JD, Erlich HA, et al. Genome-wide association study and meta-analysis find that over 40 loci affect risk of type 1 diabetes. Nature genetics. 2009;41(6):703–7.

13. Kiani AK, John P, Bhatti A, Zia A, Shahid G, Akhtar P, et al. Association of 32 type 1 diabetes risk loci in Pakistani patients. Diabetes research and clinical practice. 2015;108(1):137–42.

14. Reddy MV, Wang H, Liu S, Bode B, Reed JC, Steed RD, et al. Association between type 1 diabetes and GWAS SNPs in the southeast US Caucasian population. Genes and immunity. 2011;12(3):208–12.

15. Smith MJ, Pastor L, Newman JRB, Concannon P. Genetic Control of Splicing at SIRPG Modulates Risk of Type 1 Diabetes. Diabetes. 2022;71(2):350–8.

16. O’Connor D, Png E, Khor CC, Snape MD, Hill AVS, van der Klis F, et al. Common Genetic Variations Associated with the Persistence of Immunity following Childhood Immunization. Cell reports. 2019;27(11):3241–53 e4.

17. Sinha S, Borcherding N, Renavikar PS, Crawford MP, Tsalikian E, Tansey M, et al. An autoimmune disease risk SNP, rs2281808, in SIRPG is associated with reduced expression of SIRPgamma and heightened effector state in human CD8 T-cells. Sci Rep. 2018;8(1):15440.

18. Sinha S, Renavikar PS, Crawford MP, Steward-Tharp SM, Brate A, Tsalikian E, et al. Altered expression of SIRPgamma on the T-cells of relapsing remitting multiple sclerosis and type 1 diabetes patients could potentiate effector responses from T-cells. PLoS One. 2020;15(8):e0238070.

19. Dehmani S, Nerriere-Daguin V, Neel M, Elain-Duret N, Heslan JM, Belarif L, et al. SIRPgamma-CD47 Interaction Positively Regulates the Activation of Human T Cells in Situation of Chronic Stimulation. Front Immunol. 2021;12:732530.

20. Mao G, Li J, Wang N, Yu H, Han S, Xiang M, et al. SIRPG promotes lung squamous cell carcinoma pathogenesis via M1 macrophages: a multi-omics study integrating data and Mendelian randomization. Front Oncol. 2024;14:1392417.

21. Xu C, Jin G, Wu H, Cui W, Wang YH, Manne RK, et al. SIRPgamma-expressing cancer stem-like cells promote immune escape of lung cancer via Hippo signaling. J Clin Invest. 2022;132(5).

22. Sharp RC, Brown ME, Shapiro MR, Posgai AL, Brusko TM. The Immunoregulatory Role of the Signal Regulatory Protein Family and CD47 Signaling Pathway in Type 1 Diabetes. Front Immunol. 2021;12:739048.

23. Ye X, Wang X, Lu R, Zhang J, Chen X, Zhou G. CD47 as a potential prognostic marker for oral leukoplakia and oral squamous cell carcinoma. Oncol Lett. 2018;15(6):9075–80.

24. Azzam HS, DeJarnette JB, Huang K, Emmons R, Park CS, Sommers CL, et al. Fine tuning of TCR signaling by CD5. J Immunol. 2001;166(9):5464–72.

25. Liu X, Wang Y, Lu H, Li J, Yan X, Xiao M, et al. Genome-wide analysis identifies NR4A1 as a key mediator of T cell dysfunction. Nature. 2019;567(7749):525–9.

26. Kallies A. Distinct regulation of effector and memory T-cell differentiation. Immunol Cell Biol. 2008;86(4):325–32.

27. Pritchard GH, Phan AT, Christian DA, Blain TJ, Fang Q, Johnson J, et al. Early T-bet promotes LFA1 upregulation required for CD8+ effector and memory T cell development. J Exp Med. 2023;220(2).

28. Stefanidakis M, Newton G, Lee WY, Parkos CA, Luscinskas FW. Endothelial CD47 interaction with SIRPgamma is required for human T-cell transendothelial migration under shear flow conditions in vitro. Blood. 2008;112(4):1280–9.

29. Takahashi S. Molecular functions of SIRPalpha and its role in cancer. Biomed Rep. 2018;9(1):3–7.

