## Supplementary figures and images for "SIRPγ modulates effector differentiation of human CD8 T Cells under suboptimal TCR stimulation: implications for immune homeostasis and autoimmunity"

### Supplemental Fig 1

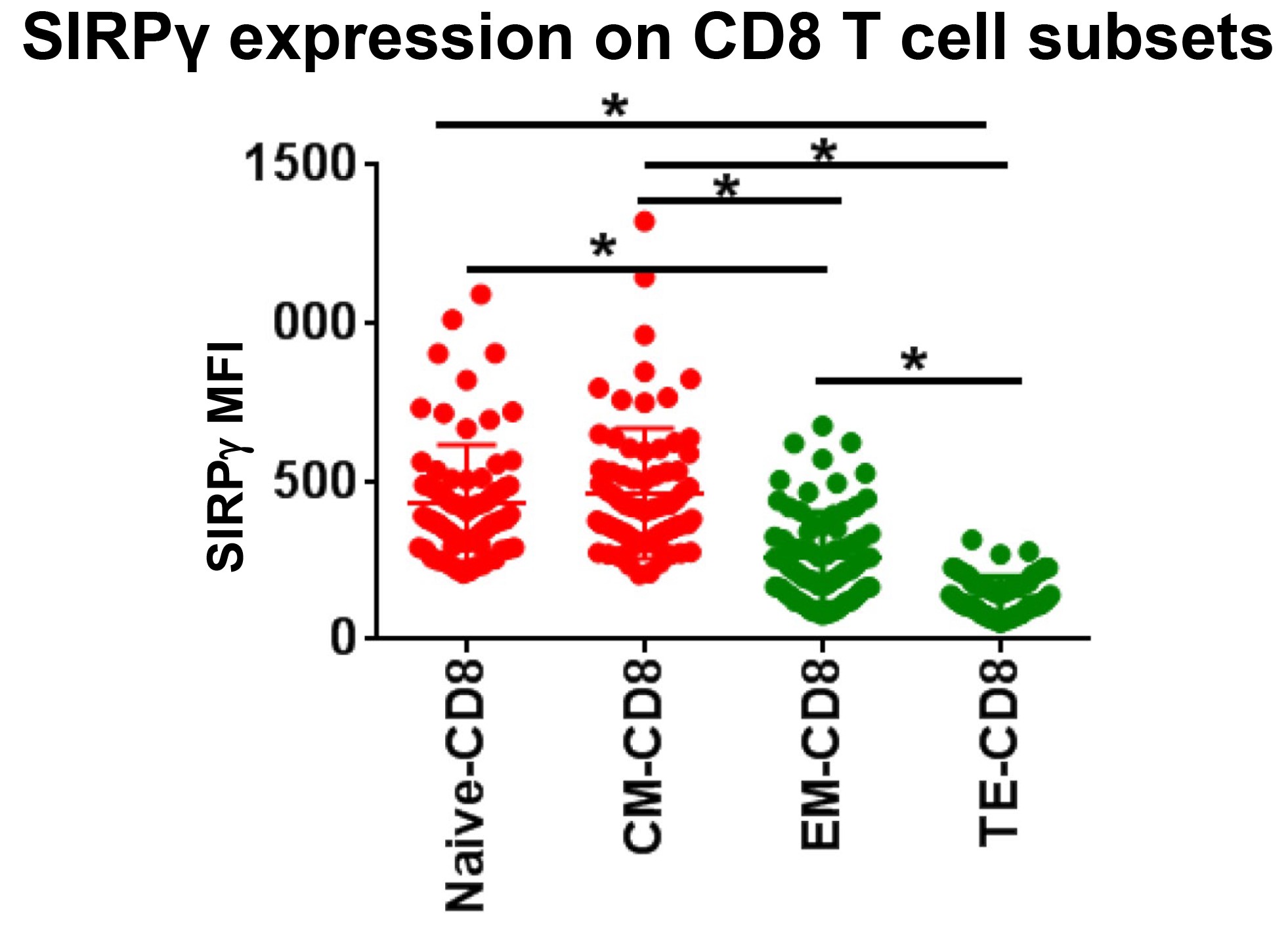
